# MakeHub: Fully automated generation of UCSC Genome Browser Assembly Hubs

**DOI:** 10.1101/550145

**Authors:** Katharina J. Hoff

## Abstract

Novel genomes are today often annotated by small consortia or individuals whose background is not from bioinformatics. This audience requires tools that are easy to use. This need had been addressed by several genome annotation tools and pipelines. Visualizing resulting annotation is a crucial step of quality control. The UCSC Genome Browser is a powerful and popular genome visualization tool. Assembly Hubs allow browsing genomes that are hosted locally via already available UCSC Genome Browser servers. The steps for creating custom Assembly Hubs are well documented and the required tools are publicly available. However, the number of steps for creating a novel Assembly Hub is large. In some cases the format of input files needs to be adapted which is a difficult task for scientists without programming background. Here, we describe the novel command line tool MakeHub that generates Assembly Hubs for the UCSC Genome Browser in a fully automated fashion. The pipeline also allows extending previously created Hubs by additional tracks.

MakeHub is freely available for download from https://github.com/Gaius-Augustus/MakeHub.

## 1 Introduction

With decreasing sequencing costs, sequencing the genomes of non-model organisms that are of interest to individuals or small research consortia has become affordable. Pipelines and tools that enable scientists with diverse backgrounds to easily annotate protein coding genes in novel genomes have been developed and are frequently used, for example the tools AUGUSTUS (Stanke *et al.*, 2008), GeneMark-ES/ET (Lomsadze *et al*., 2014) and GeMoMa (Keilwagen *et al.*, 2018), and the pipelines BRAKER (Hoff *et al.*, 2015), WebAUGUSTUS (Hoff and Stanke, 2013) and MAKER (Holt and Yandell, 2011). The output of such gene prediction tools and pipelines is in a table-like text file GTF or GFF3 format. Visualization of predicted gene structures in context with available extrinsic evidence is a crucial step of quality control in any genome annotation project (Hoff and Stanke, 2015). A number of genome browsers are available for this task, for example the UCSC Genome Browser (Kent *et al.*, 2002), JBrowse (Skinner *et al.*, 2009) and GBrowse2 (Stein, 2013). While JBrowse and GBrowse2 require installation of the browser software on a server or on a local computer, the UCSC Genome Browser bypasses the requirement for installation by offering the opportunity of visualizing any genome through so-called locally hosted ‘Assembly Hubs’ combined with already existing UCSC Genome Browser servers, e.g. at https://genome.ucsc.edu (February 10^*th*^ 2019) (Raney *et al.*, 2013).

An Assembly Hub is simply a directory that contains configuration files required by the UCSC Genome Browser as well as track data files with the data to be visualized. The steps for creating custom Assembly Hubs are well documented (for example at http://genomewiki.ucsc.edu/index.php/Assembly_Hubs, February 10^*th*^ 2019) and the required tools are publicly available. An experienced bioinformatician will be able to create Assembly Hubs with ease. However, a scientist with limited programming background will find it troublesome to manually create the required configuration files, to adapt the output of gene prediction pipelines to the demands of UCSC tools for creating data tracks, and to run all required tools in the correct order. Recently, a Galaxy workflow for fully automated generation of UCSC Assembly Hubs (and JBrowse instances) became available (Liu *et al.*, 2018). But not all groups run their own Galaxy instance. Therefore, we here describe the novel command line tool MakeHub for the fully automated generation of UCSC Assembly Hubs from the output of BRAKER, MAKER and GeMoMa for a genome of single species.

## 2 MakeHub pipeline

The MakeHub pipeline is implemented in Python and is compatible with Linux and Mac OS X x86 64 computers. The pipeline is illustrated in figure 1. It provides a command line interface for creating fully functional Assembly Hubs from a genome file (and optionally files with gene and evidence information) and for adding tracks to existing hubs.

### 2.1 Genome

Genome files in FASTA format are converted to 2bit format using UCSC’s `faToTwoBit`. Chromosome (or contig) sizes that serve as input to tools for the later creation of bigWig and bigBed files are extracted using UCSC’s `twoBitInfo`. GC-content information is collected from the genome with UCSC’s `hgGcPercent` and written into WIG format. Subsequently, the WIG file is converted to bigWig format using UCSC’s `wigToBigWig`. The genome is screened for softmasked repeat information by MakeHub and repeats are written to BED format, sorted and then converted to bigBed format with UCSCs `bedToBigBed`.

Hub configuration files, most prominently the `hub/hub.txt` file, are created and initialized upon completion of genome processing.

### 2.2 RNA-Seq

Transcriptome alignment files in BAM format can be visualized in two ways. By default, BAM-files are sorted with SAMtools (Heng *et al.*, 2009) and converted to WIG format. This conversion step can either be performed with the AUGUSTUS tool bam2wig (Hoff and Stanke, 2018) or, in the absence of that tool, with SAMtools and built-in MakeHub functionality. WIG-files are subsequently converted to bigWig format as described above. This generates tracks that allow for an intuitive interpretation of gene structures in context with RNA-Seq coverage information.

Optionally, BAM-files can be displayed from native BAM format. The required SAM index is automatically generated with SAMtools. Viewing native BAM-files gives immediate access to alignment quality information of single reads.

### 2.3 Gene Models

MakeHub seamlessly integrates with output files of the popular genome annotation tools and pipelines AUGUSTUS, GeneMark-ES/ET, BRAKER, MAKER and GeMoMa. GTF- and GFF3-files of these tools and pipelines are standardized to a UCSC-compatible GTF format by MakeHub. Sub-sequently, UCSC’s `gtfToGenePred` is used to convert to GenePred data, which is checked for consistency by UCSC’s `genePredCheck` and passed to `bedToBigBed` after sorting, generating track data in bigBed format.

**Figure 1:**
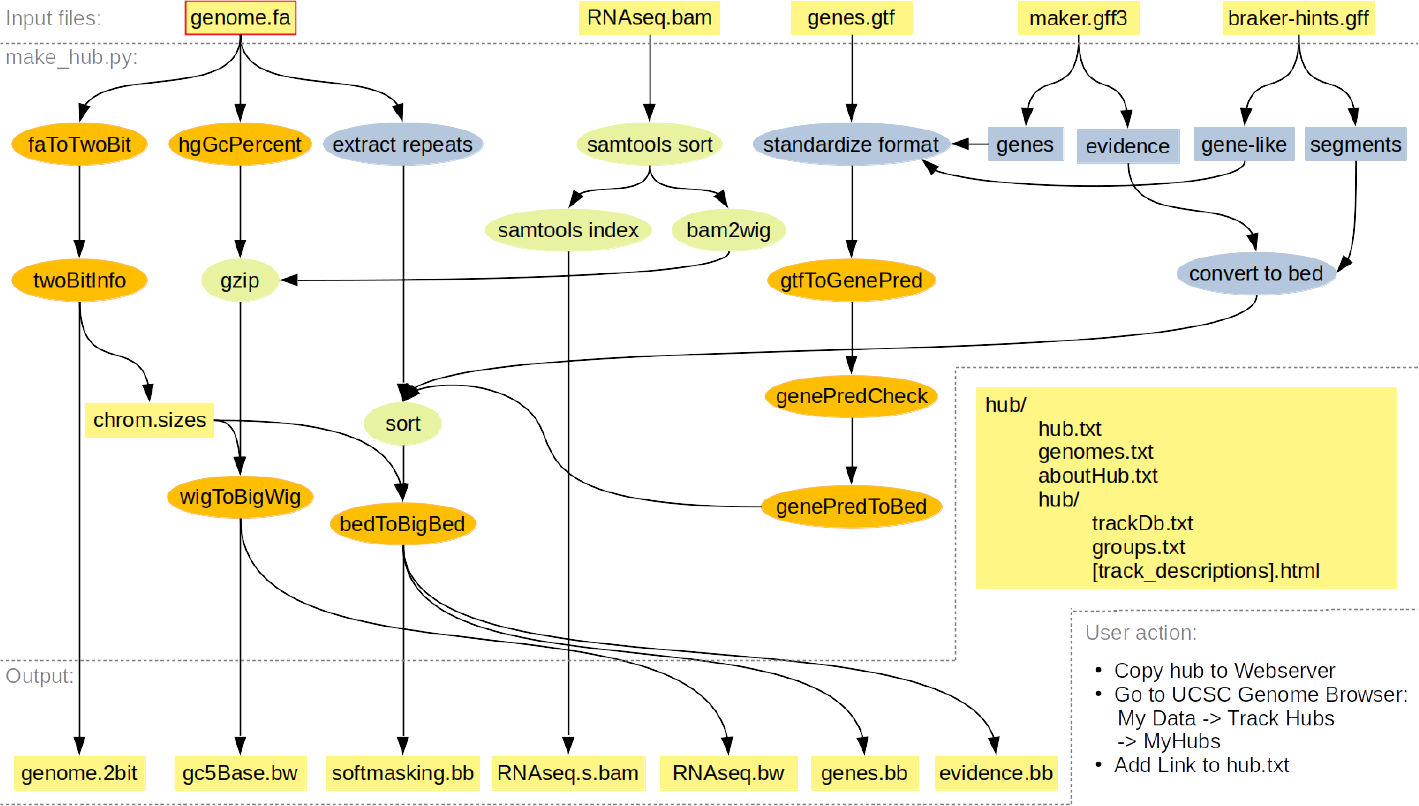
Illustration of the MakeHub pipeline. Input and output files and directories are shown in yellow (the only compulsory input file for creating a new hub is makred with a red frame, all other input files are optional), UCSC tools are labeled with orange color. Other external tools are labeled in green, MakeHub components are labeled in blue.

MakeHub accepts the output directory of a BRAKER run as an input argument and automatically identifies the gene prediction files for visualization in that directory (alternatively, AUGUSTUS and GeneMark-ES/ET predictions can be passed as arguments to separate options). MakeHub automatically extracts gene models from the MAKER GFF3 output file (it usually contains evidence for gene models as well). MakeHub accepts the native GFF3 output file of GeMoMa.

### 2.4 Evidence

Visualization of the evidence that went into gene model inference is crucial. This evidence often exceeds the information that can be seen in a RNA-Seq WIG or BAM track. For example, annotators are often interested in viewing splice-junctions from RNA-Seq and/or protein alignments with coverage and strand information in a concise overview. On the other hand, alignments from cDNA, assembled transcriptomes, and proteins need to be visualized in a gene-structure-like fashion. MakeHub automatically generates suitable tracks with evidence from MAKER output and from BRAKER hints files in GFF format. Gene-structure-like evidence (e.g. full length protein alignments) are visualized similar to gene models, while other evidence, such as splice-junctions, is visualized as segments. All resulting evidence track data files are in the indexed bigBed format.

### 2.5 Hub and Track Descriptions

MakeHub automatically generates HTML template files for describing a hub and its tracks. These files are required for public hubs (http://genomewiki.ucsc.edu/index.php/Public_Hub_Guidelines, February 10^*th*^ 2019). These pages should be edited to appropriately describe genome projects and individual tracks before adding a hub to the list of public hubs at UCSC.

### 2.6 Deploying Assembly Hubs from MakeHub

Automatically generated Assembly Hubs must be copied to a publicly available web server for deployment. The hyperlink to hub.txt can be provided to the UCSC Genome Browser for data visualization (see instructions in Figure 1).

## 3 Conclusion

In summary, MakeHub is a command line tool that enables scientists with little experience in bioinformatics to generate Assembly Hubs of their genome and annotation of interest with ease.

## Acknowledgements

The international collaboration between the groups of Mark Borodovsky and Mario Stanke, supported by US National Institutes of Health grant HG000783, gave rise to the development of MakeHub.

I thank Mario Stanke, Matthis Ebel and Malte Wellnitz for proofreading.

